## Supplemental Methods for "Caspase-mediated nuclear pore complex trimming in cell differentiation and endoplasmic reticulum stress"

**Table 1. Primary antibodies used in this study**

| Antigen group | Antigen | Host | Clonality | Application | Dilution | Source |
|---|---|---|---|---|---|---|
| Nups | Tpr | Rabbit | Polyclonal | Western blot | 1:1000 | Abcam (ab84516) |
| | | | | Immunofluorescence | 1:300 | |
| | | Mouse | Monoclonal | Immunofluorescence | 1:200 | Santa Cruz (sc-271565) |
| | Nup358<br>Nup214<br>Nup153<br>Nup62 | Mouse | Monoclonal | Western blot | 1:4000 | Covance (MMS-120R) |
| | | | | Immunofluorescence | 1:500 | |
| | Nup153 | Mouse | Monoclonal | Immunofluorescence | 1:3 | In-house (SA1) |
| | | Rat | Monoclonal | Immunofluorescence | 1:300 | Abcam (ab81463) |
| | Nup96 | Rabbit | Monoclonal | Western blot | 1:1000 | Abcam (ab124980) |
| | Nup93 | Mouse | Monoclonal | Western blot | 1:200 | Santa Cruz (sc-81343) |
| | Nup50 | Rabbit | Monoclonal | Western blot | 1:1000 | Abcam (ab151567) |
| | ELYS | Rabbit | Polyclonal | Western blot | 1:1000 | In-house |
| | Nup205 | Rabbit | Polyclonal | Western blot | 1:500 | In-house |
| | Nup188 | Rabbit | Polyclonal | Western blot | 1:1000 | Bethyl Laboratories (A302-323A) |
| | Nup160 | Rabbit | Polyclonal | Western blot | 1:500 | In-house |
| | Nup155 | Guinea Pig | Polyclonal | Western blot | 1:1000 | In-house |
| | Nup85 | Rabbit | Polyclonal | Western blot | 1:500 | In-house |

| | | | | | | |
|---|---|---|---|---|---|---|
| | Sec13 | Rabbit | Polyclonal | Western blot | 1:500 | In-house |
| | Nup37 | Rabbit | Polyclonal | Western blot | 1:500 | In-house |
| | Pom121 | Rabbit | Polyclonal | Western blot | 1:1000 | Invitrogen (PA5-36498) |
| | | | | Immunofluorescence | 1:300 | |
| Caspases | Caspase-3 | Rabbit | Polyclonal | Western blot | 1:1000 | Cell Signaling Technology (9661) |
| | Caspase-9 | Rabbit | Polyclonal | Western blot | 1:1000 | Cell Signaling Technology (9504) |
| | Caspase-12 | Rat | Monoclonal | Western blot | 1:200 | Santa Cruz (sc-21747) |
| IAPs | XIAP | Rabbit | Polyclonal | Western blot | 1:1000 | Cell Signaling Technology (2042) |
| | Survivin | Rabbit | Monoclonal | Western blot | 1:1000 | Cell Signaling Technology (2808) |
| Differentiation-related | Cleaved Notch1 | Rabbit | Monoclonal | Western blot | 1:1000 | Cell Signaling Technology (4147) |
| | Myogenin | Mouse | Monoclonal | Western blot | 1:1000 | BD Biosciences (556358) |
| | | | | Immunofluorescence | 1:200 | |
| | Myosin heavy chain | Mouse | Monoclonal | Western blot | 1:100 | In-house (MF20) |
| | Sox2 | Rabbit | Polyclonal | Western blot | 1:1000 | Cell Signaling Technology (2748) |
| | βIII-Tubulin | Rabbit | Polyclonal | Western blot | 1:5000 | Biolegend (802001) |
| ER stress | Bip | Rabbit | Monoclonal | Western blot | 1:1000 | Cell Signaling Technology (3177) |
| Caspase substrates | PARP | Rabbit | Monoclonal | Western blot | 1:1000 | Cell Signaling Technology (9532) |
| | αII-Spectrin | Mouse | Monoclonal | Western blot | 1:100 | Santa Cruz (sc-48382) |
| Cytoplasm & | α-Tubulin | Mouse | Monoclonal | Western blot | 1:5000 | Sigma (T5168) |

| | | | | | | |
|---|---|---|---|---|---|---|
| nucleoplasm markers | Lamin B1 | Mouse | Monoclonal | Western blot | 1:200 | Santa Cruz (sc-374015) |
| | Fibrillarin | Chicken | Polyclonal | Western blot | 1:2000 | Novus Biologicals (NBP2-46881) |
| Focal adhesion proteins | Hic-5 | Rabbit | Polyclonal | Western blot | 1:1000 | Proteintech (10565-1-AP) |
| | Zyxin | Mouse | Monoclonal | Western blot | 1:1000 | R&D Systems (MAB6977) |
| | Paxillin | Mouse | Monoclonal | Western blot | 1:1000 | Invitrogen (AHO0492) |
| | FAK | Mouse | Monoclonal | Western blot | 1:1000 | BD Biosciences (610087) |
| | | Rabbit | Polyclonal | Western blot | 1:1000 | Cell Signaling Technology (3285) |
| Karyopherins | Crm1 | Mouse | Monoclonal | Immunofluorescence | 1:200 | BD Biosciences (611833) |
| | Importin-α | Mouse | Monoclonal | Immunofluorescence | 1:1000 | Abcam (ab2811) |
| | Importin-β | Rabbit | Polycloncal | Immunofluorescence | 1:200 | Novus Biologicals (NBP2-38482) |
| RNA polymerase | Phospho-Rbp1 (Ser5) | Rabbit | Monoclonal | Western blot | 1:1000 | Cell Signaling Technology (13523) |

**Table 2. Secondary antibodies used in this study**

| Antibody | Host | Clonality | Application | Dilution | Source |
|---|---|---|---|---|---|
| HRP-conjugated anti-mouse IgG | Goat | Polyclonal | Western blot | 1:10,000 | Invitrogen (G-21040) |
| HRP-conjugated anti-rabbit IgG | Goat | | | 1:10,000 | Invitrogen (G-21234) |
| HRP-conjugated anti-rat IgG | Goat | | | 1:3,000 | Invitrogen (A10549) |
| HRP-conjugated anti-chicken IgY | Goat | | | 1:3000 | Invitrogen (A16054) |
| HRP-conjugated anti-guinea pig IgG | Goal | | | 1:3000 | Invitrogen (A18775) |
| IRDye800-conjugated anti-mouse IgG | Donkey | | | 1:10,000 | Rockland Immunochemicals (610-732-124) |
| Alexa Fluor 680-conjugated Anti-rabbit IgG | Goat | | | 1:10,000 | Invitrogen (A-21109) |
| Alexa Fluor 647-conjugated anti-mouse IgG | Donkey | | Immunofluorescence | 1:1000 | Invitrogen (A-31571) |
| Alexa Fluor 568-conjugated Anti-rabbit IgG | Goat | | | 1:1000 | Invitrogen (A-11036) |
| Alexa Fluor 488-conjugated anti-rat IgG | Goat | | | 1:1000 | Invitrogen (A-11006) |

**Table 3. Chemicals used in this study**

| Chemical | Function | Stock solution solvent | Stock solution concentration | Working concentration | Source |
|---|---|---|---|---|---|
| Q-VD(OMe)-OPh | Pan-caspase inhibitor | DMSO | 30 mM | 30 μM | APExBIO Technology (A8165) |
| Z-LL-CHO | Pan-calpain inhibitor | | 50 mM | 50 μM | Peptide Institute, Inc. (IZL-3178-v) |
| FK506 | Calcineurin inhibitor | | 1 mg/ml | 0-100 ng/ml | Enzo Life Sciences (ALX-380-008) |
| DAPT | γ-Secretase inhibitor | | 5 mM | 0-10 μM | Enzo Life Sciences (ALX-270-416) |
| Tunicamycin | ER stress inducer | | 10 mg/ml | 0-1 μg/ml | Tocris (3516) |
| SCH772984 | ERK1/2 inhibitor | | 10 mM | 1 μM | BioVision (B1682-5) |
| Doxycycline | Tetracycline transactivator activator | $H_2O$ | 1 mg/ml | 0-1000 ng/ml | Alfa Aesar (J60422) |
| Leptomycin B | Exportin-1 inhibitor | ethanol | 250 μM | 0-25 nM | BioVision (1814) |

### Table 4. Plasmids used in this study

| No. | Plasmid | Backbone | Insert | Insert PCR template | Insert PCR Primers (5’ to 3’) | Cloning method |
|---|---|---|---|---|---|---|
| 1 | Doxycycline-inducible GFP-myogenin | LT3GEPIR (Addgene 111177) | Myogenin | GE Healthcare Dharmacon, Inc. MMM1013-202805422 | Forward: atcgtgtacaagtcatcagagctgtatgagacatccccctatttc<br>Reverse: atcggaattctcagttgggcatggtttcgtc | Backbone and insert cut with BsrG1 and EcoR1 were ligated by T7 DNA ligase. |
| 2 | Doxycycline-inducible GFP-myogenin | | GFP-myogenin | Plasmid No. 1 | Forward: aagtcgagcttgcgttggatc<br>Reverse: aaggcacagtgtacatcagttgggcatggtttcgtc | PCR products were cloned into the backbone cut with BamH1 and EcoR1 by In-fusion cloning. |
| | | | Bovine growth hormone Poly-A | N/A | Forward: actgatgtacactgtgccttctagttgcc<br>Reverse: acaagataattgctcgaattcccatagagcccaccgcatc | |
| 3 | EF1α promoter NES-eGFP | pEF1α-Tet3G | $NES_{Rev}$-GFP -IRES-Puro$^r$ | Plasmid DNA and map available upon request | | |
| 4 | | | $NES_{PKI}$-GFP -IRES-Puro$^r$ | | | |

$NES_{Rev}$: LPPLERLTL
$NES_{PKI}$: LALKLAGLDI

### Table 5. RNA FISH probes used in the study

| RNA target | Label | Source | Probe type | Sequence used to generate Stellaris custom probe sets (NCBI reference sequence) |
|---|---|---|---|---|
| 18S rRNA | Quasar 570 | Stellaris | Deoxyribonucleic acid | NR_003278.3[1] |
| *Gapdh* mRNA | Quasar 670 | | | NM_008084.2 (ShipReady, SMF-3140-1) |
| Poly-A RNA | TYE563 | Qiagen | Locked nucleic acid ($T_{25}$) | |
