## Supplemental Figures for "Caspase-mediated nuclear pore complex trimming in cell differentiation and endoplasmic reticulum stress"

### Figure S1 (related to Figure 1)

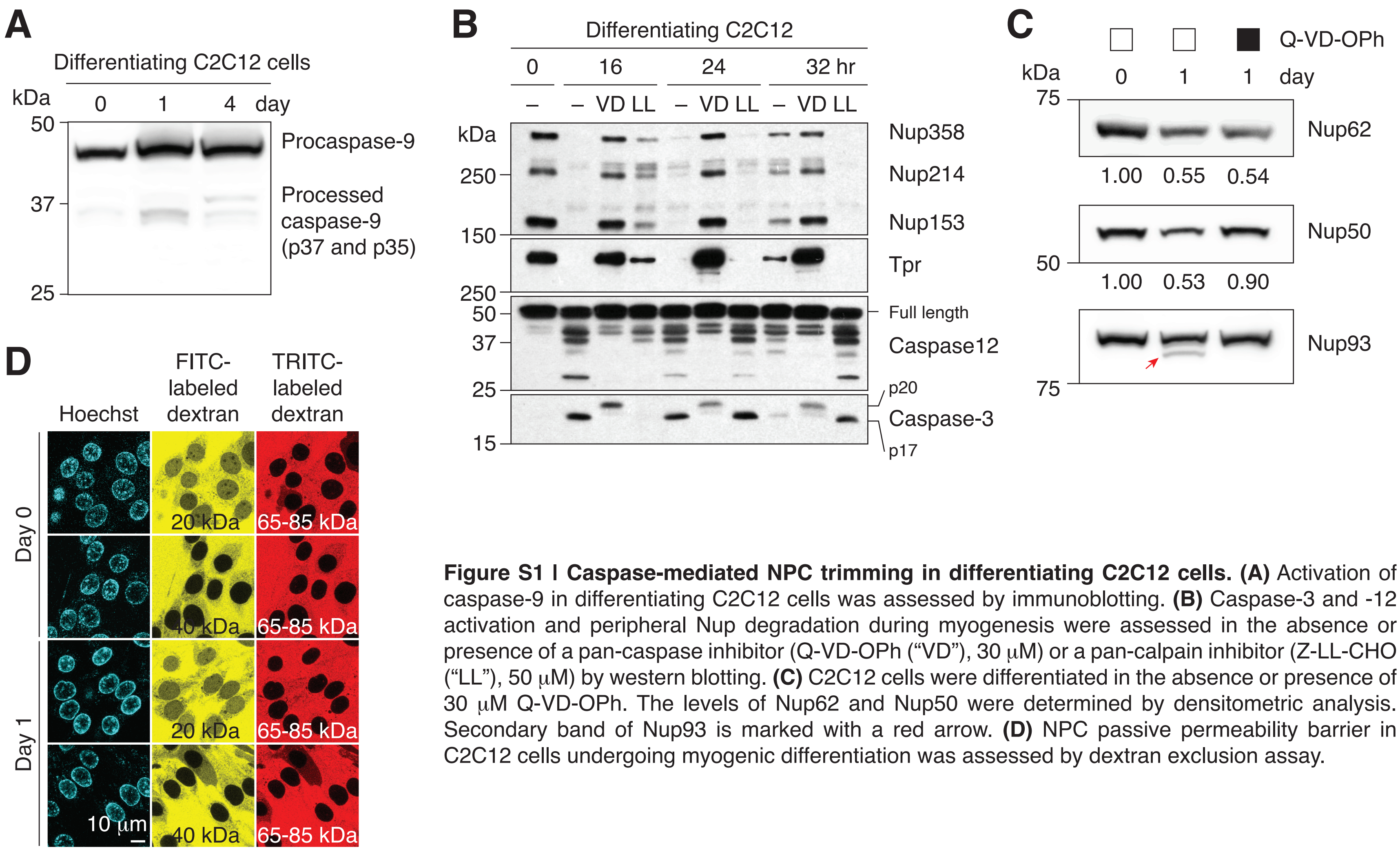


**Figure S1 | Caspase-mediated NPC trimming in differentiating C2C12 cells. (A)** Activation of caspase-9 in differentiating C2C12 cells was assessed by immunoblotting. **(B)** Caspase-3 and -12 activation and peripheral Nup degradation during myogenesis were assessed in the absence or presence of a pan-caspase inhibitor (Q-VD-OPh (“VD”), 30 μM) or a pan-calpain inhibitor (Z-LL-CHO (“LL”), 50 μM) by western blotting. **(C)** C2C12 cells were differentiated in the absence or presence of 30 μM Q-VD-OPh. The levels of Nup62 and Nup50 were determined by densitometric analysis. Secondary band of Nup93 is marked with a red arrow. **(D)** NPC passive permeability barrier in C2C12 cells undergoing myogenic differentiation was assessed by dextran exclusion assay.

### Figure S2 (related to Figure 1)

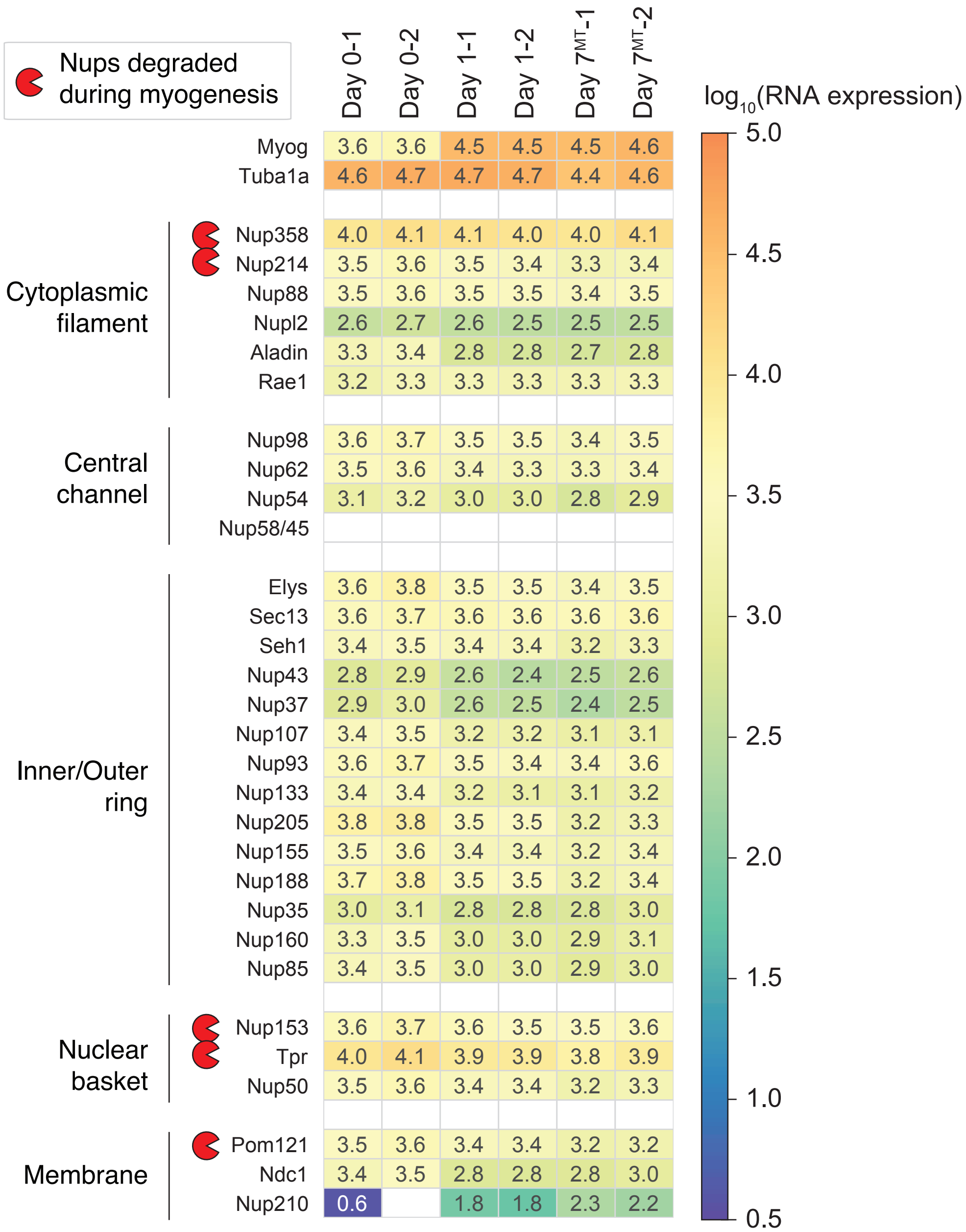


**Figure S2 | Transcript levels of NPC subunits in differentiating C2C12 cells.** Using RNA-seq, we monitored how the transcript levels of 30 Nups change over the course of myogenesis. Those of *Myog* and *Tuba1a* are also shown for comparison. GEO accession number: GSE183521.

### Figure S3 (related to Figure 1)

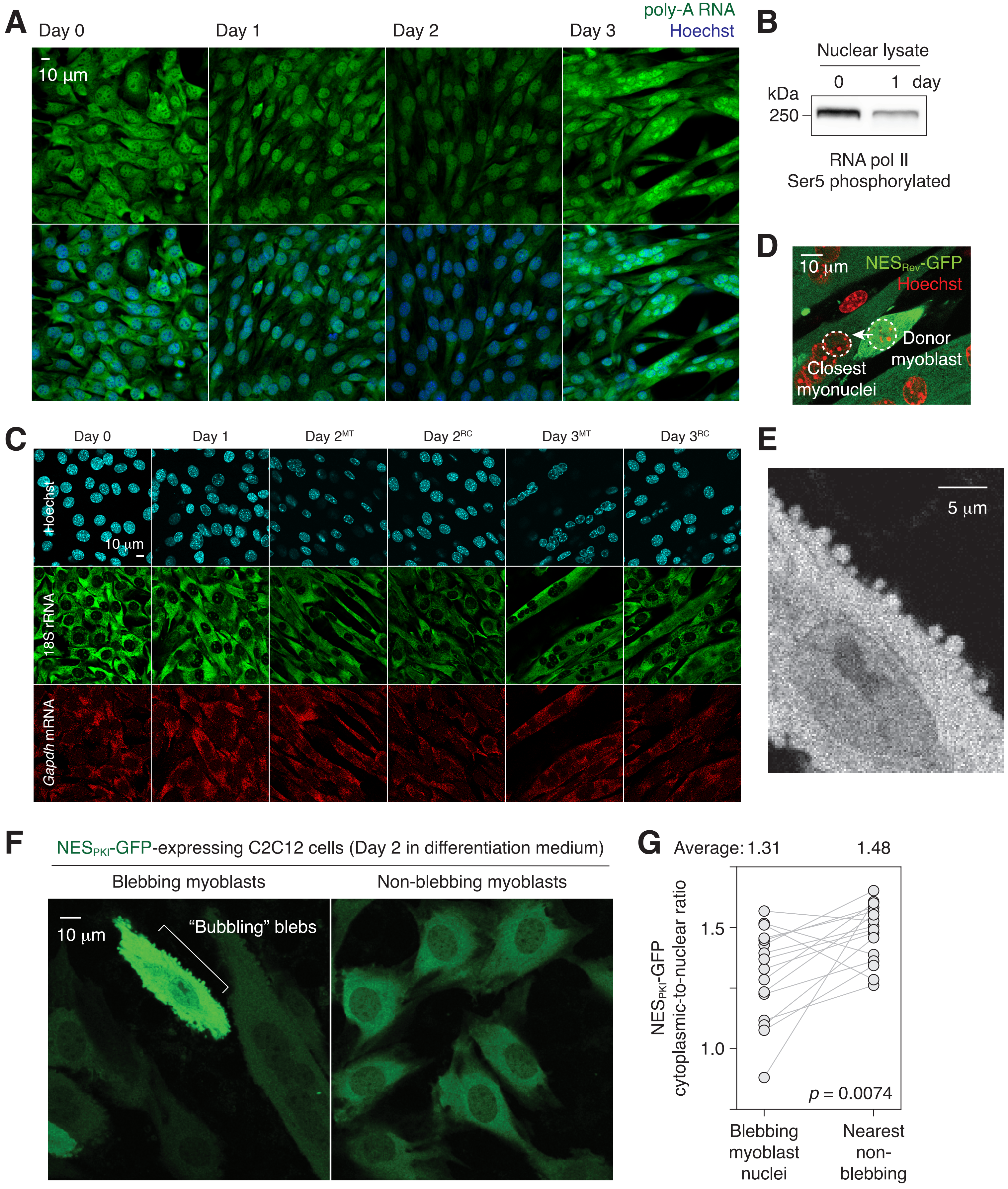

**Figure S3 l Nuclear export impairment in differentiating C2C12 cells. (A)** poly-A RNAs were visualized in differentiating C2C12 cells by fluorescence *in situ* hybridization. **(B)** Ser5 phosphorylation level of RNA polymerase II in day 0 and 1 C2C12 cells. **(C)** 18S rRNA and *Gapdh* mRNA fluorescence *in situ* images of differentiating C2C12 cells. MT: myotubes, RC: reserve cells. **(D)** Representatitive image showing how "donor myoblast" and "closest myonuclei" were determined. **(E)** Bubbling blebs on the plasma membrane of a fusion-competent myoblast. **(F)** Live imaging of C2C12 cells stably expressing $NES_{PKI}$-GFP. Imaged 48 hours after switching from growth to differentiation medium. **(G)** Cytoplasmic-to-nuclear ratio of $NES_{PKI}$-GFP was quantified for 16 pairs of blebbling and non-blebbing cells. Grey lines link each pair.

### Figure S4 (related to Figure 1)

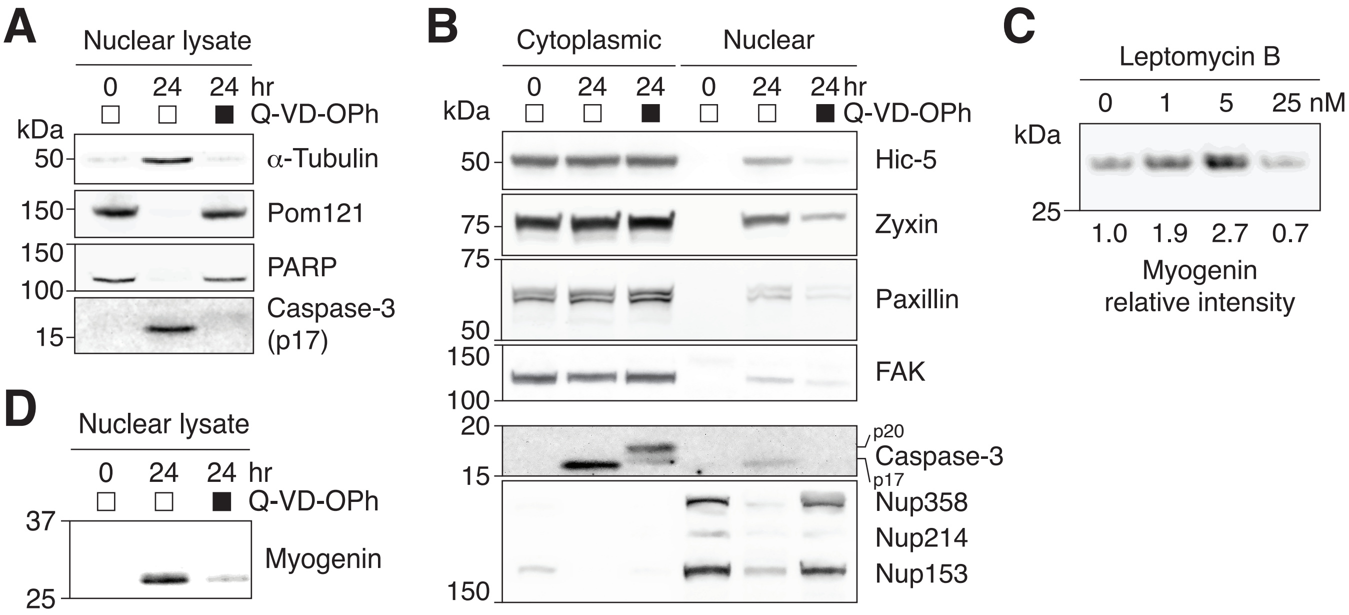


**Figure S4 l Nuclear retention of NES-containing proteins in differentiating C2C12 cells. (A)** Nuclear translocation of α-tubulin and **(D)** myogenin upregulation were assessed in C2C12 cells differentiated in the absense or presence of a pan-caspase inhibitor, Q-VD-OPh (30 μM). **(B)** C2C12 cells were differentiated in the absence or presence of Q-VD-OPh (30 μM) for 24 hours, and nuclear and cytoplasmic fractions were obtained. Nuclear accumulation of NES-containing focal adhesion proteins and caspase-mediated NPC trimming were examined by western blotting. 17.5 μg of cytoplasmic or nuclear protein loaded per lane. **(C)** C2C12 cells were differentiated in the absence or presence of leptomycin b for 24 hours, and myogenin expression was determined by western blotting.

### **Figure S5** (related to Figure 1)

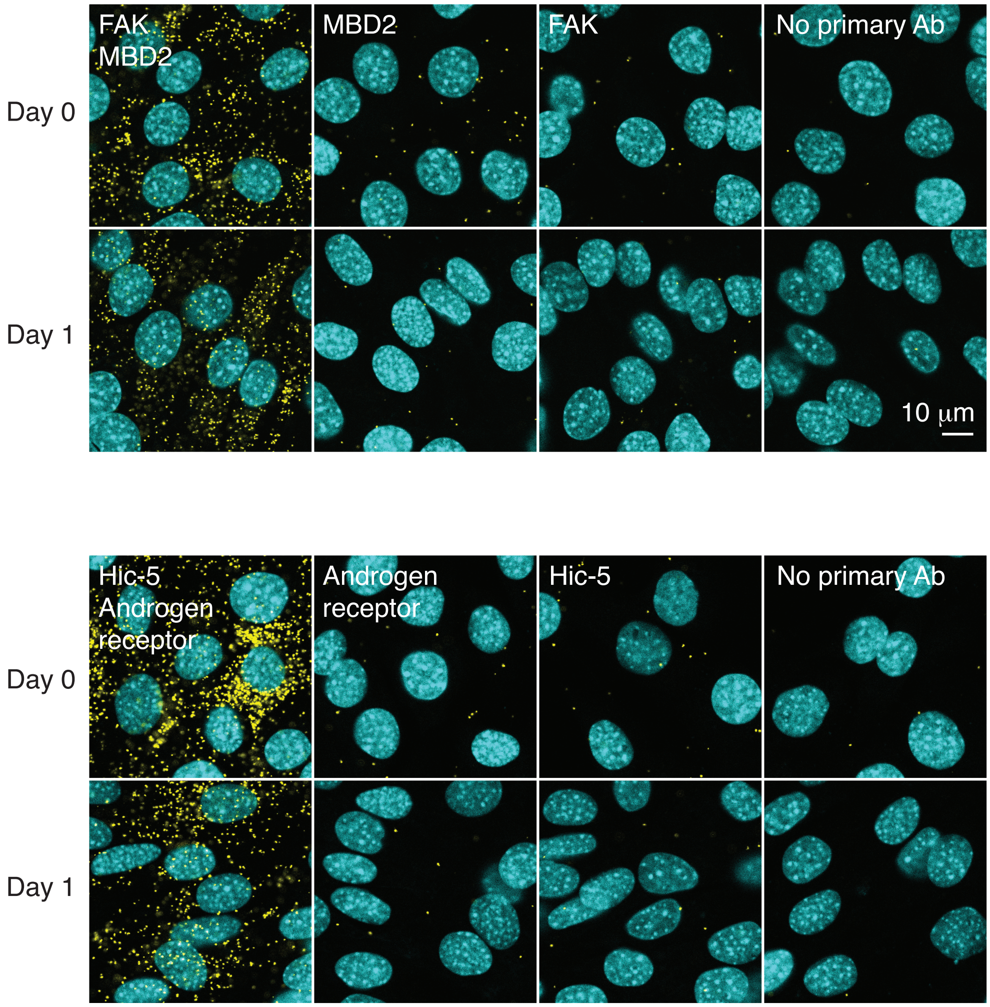


**Figure S5 I Proximity ligation assay.** Interaction between FAK and MBD2 and between Hic-5 and androgen receptor was assessed by proximity ligation assay (yellow puncta) in differentiating C2C12 cells.

### **Figure S6** (related to Figure 2)

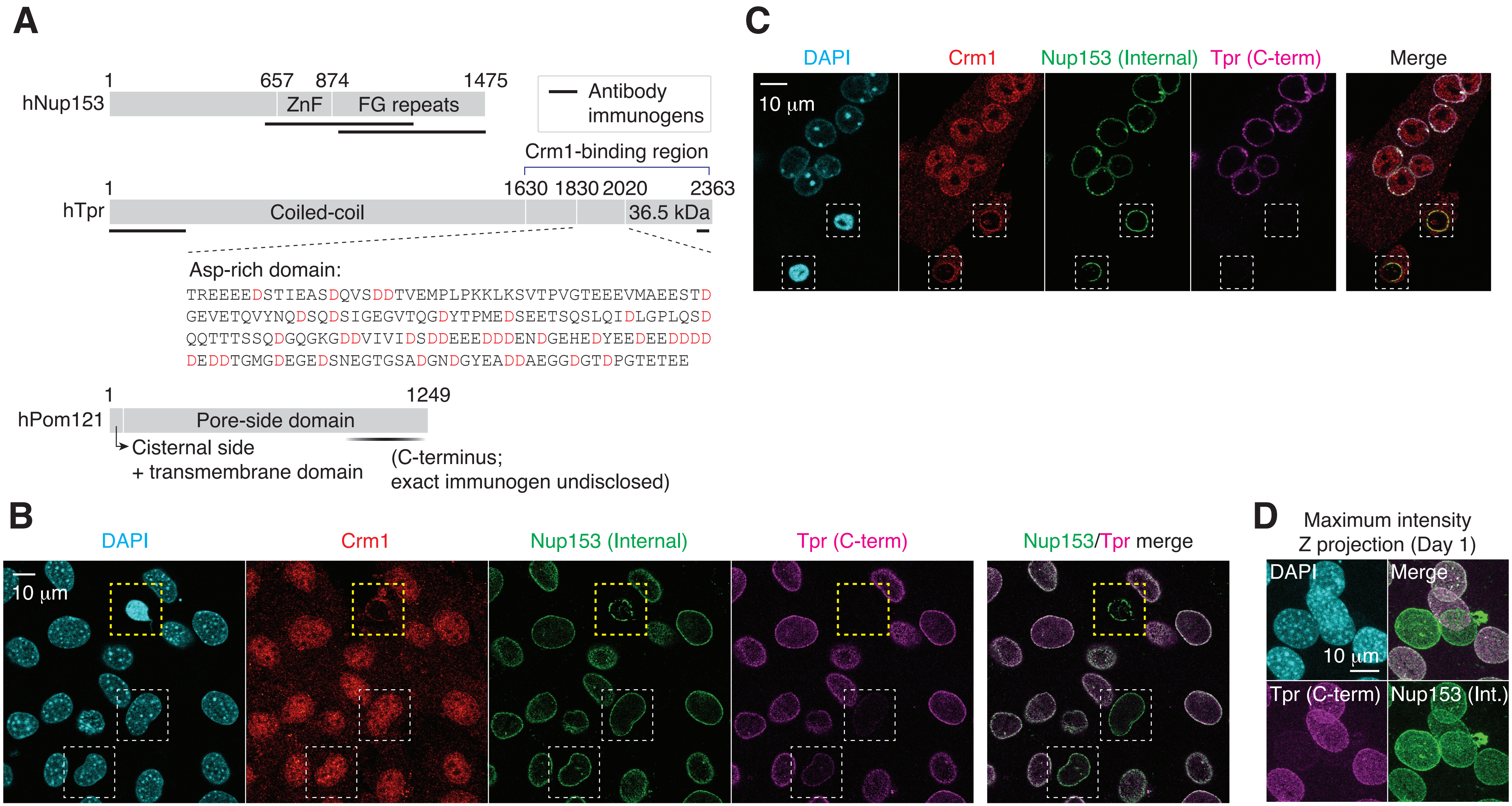


**Figure S6 I Immunofluorescence staining of apoptotic and differentiating C2C12 cells. (A)** Domain architecture of human Nup153, Tpr, and Pom121. Immunogenic fragments used to generate Nup153, Tpr, and Pom121 antibodies are mapped on each protein. Aspartates (D) in Tpr aspartate-rich region are colored in red. **(B)** C2C12 cells that have undergone myogenic differentiation for a day. In yellow dashed box is an apoptotic cell, and in white dashed boxes, differentiating TprC⁻ cells. **(C)** C2C12 cells were immunostained for Crm1, Nup153, and Tpr. Apoptotic nuclei with condensed chromatin are shown in dashed boxes. **(D)** TprC⁻ cells were identified by immunofluorescence. Maximum intensity Z projection micrographs were reconstructed from confocal imaging.

### **Figure S7** (related to Figure 2)

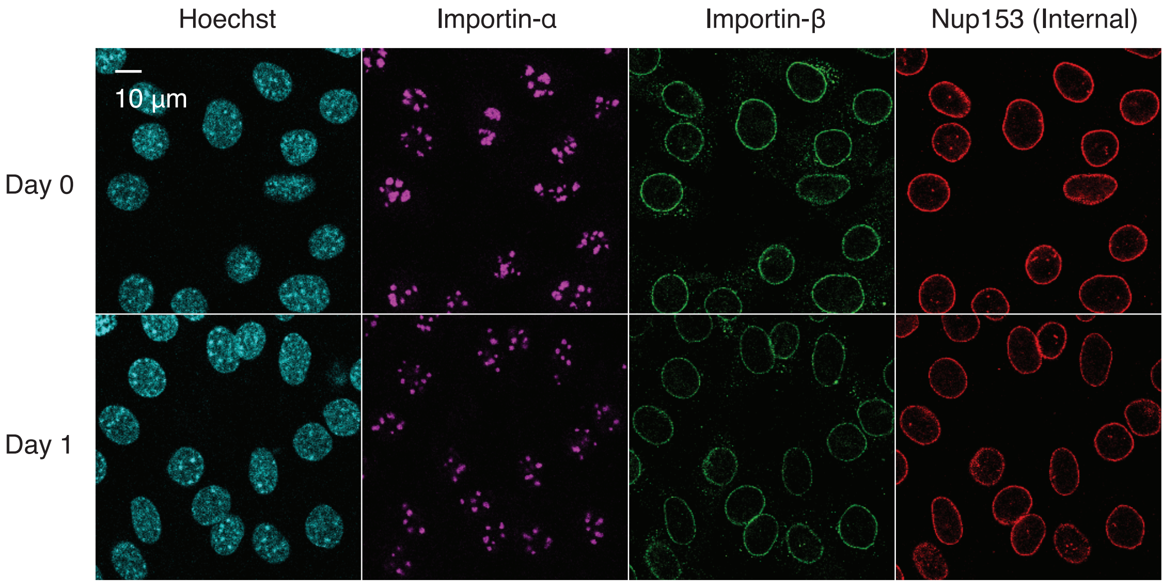


**Figure S7 l Immunofluorescence staining of differentiating C2C12 cells.** C2C12 cells undergoing myogenesis were immunostained for importin-α, importin-β, and Nup153.

#### Figure S8 (related to Figure 5)

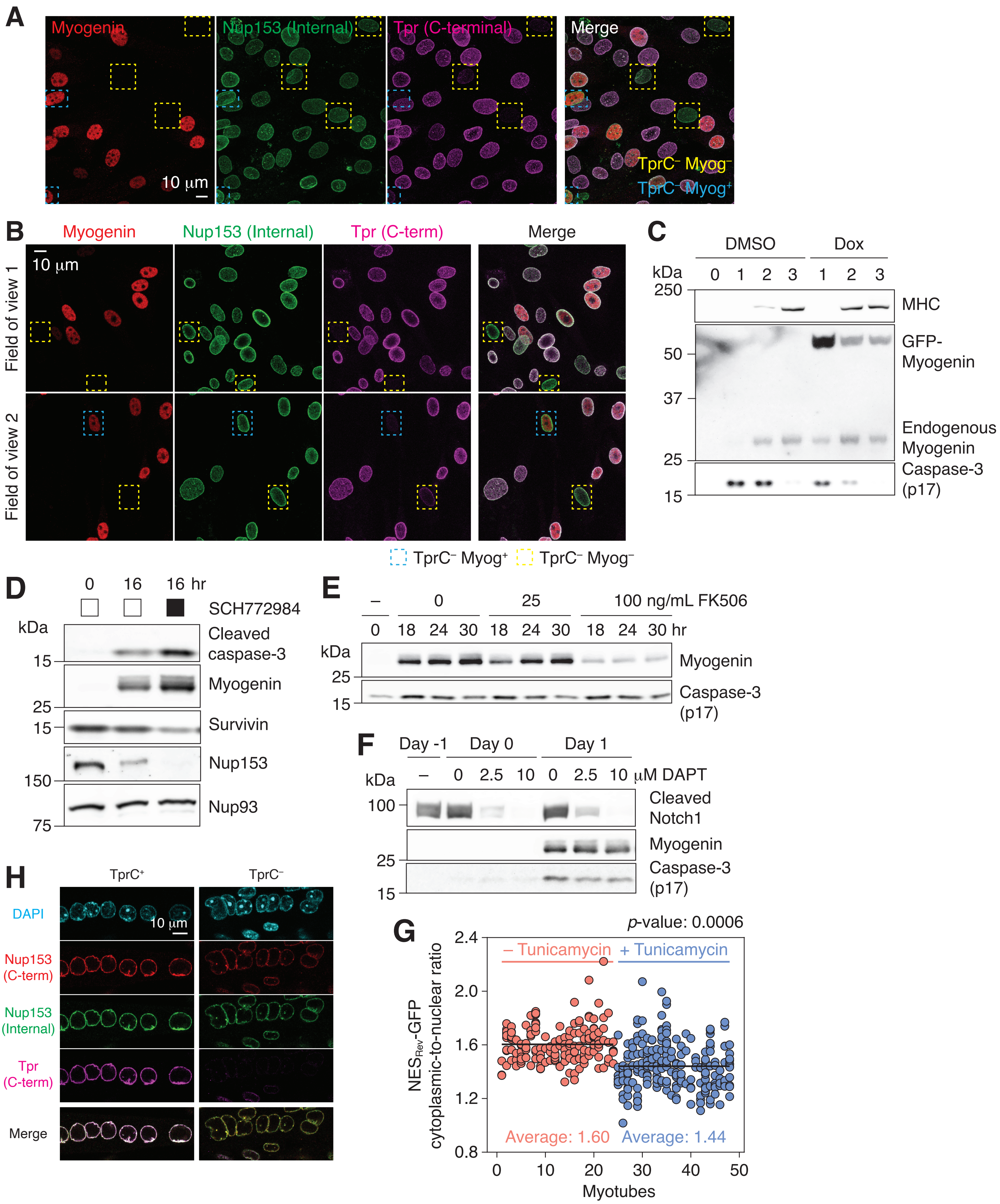

**Figure S8 | Activation of caspases and proteolysis of their substrates. (A)** C2C12 cells that have undergone myogenic differentiation for 1 day were immunostained for myogenin, Nup153, and Tpr. In yellow and blue dashed boxes are myogenin-negative and -positive TprC$^{-}$ cells, respectively. **(B)** Primary myoblasts were differentiated for 24 hours. Cells were immunostained for myogenin, Nup153, and Tpr. Two different fields of view are shown. Nuclei that lack Tpr C-terminal epiptope are boxed in yellow (myogenin-negative) or blue (myogenin-positive). **(C)** C2C12 stable cell line that expresses GFP-myogenin in a doxycycline-dependent manner was differentiated in the absence or presence of 0.5 μM doxycycline (Dox). Myosin heavy chain (MHC), exogenous and endogenous myogenin, and active caspase-3 levels were determined by immunoblotting. **(D)** C2C12 cells were differentiated in the absence or presence of an ERK1/2 inhibitor (SCH772984, 1 μM). Expression levels of active caspase-3, myogenin, survivin, Nup153, and Nup93 were determined by immunoblotting. **(E)** Upregulation of myogenin and formation of caspase-3 p17 were evaluated in the presence of 0, 25, or 100 ng/mL FK506 by immunoblotting. **(F)** Immunoblots showing the expression levels of cleaved Notch1, myogenin, and active caspase-3 in C2C12 cells. DAPT was added at 0, 2.5, or 10 μM on day -1, and maintained throughout differentiation. **(G)** The cytoplasmic-to-nuclear ratio of NES-GFP was determined for myonuclei in 24 myotubes treated with DMSO (red) or tunicamycin (blue, 1 μg/mL for 24 hours). Each data point represents a myonucleus, and ones that share the same x values are from the same myotube. At least 3 myonuclei per myotube were quantified. **(H)** Myotubes were treated with 1 μg/mL tunicamycin for 24 hours, and immunostained for Nup153 and Tpr. Tpr C-terminal epitope was undetectable (TprC$^{-}$) in 5-10% of the population.

### Figure S9

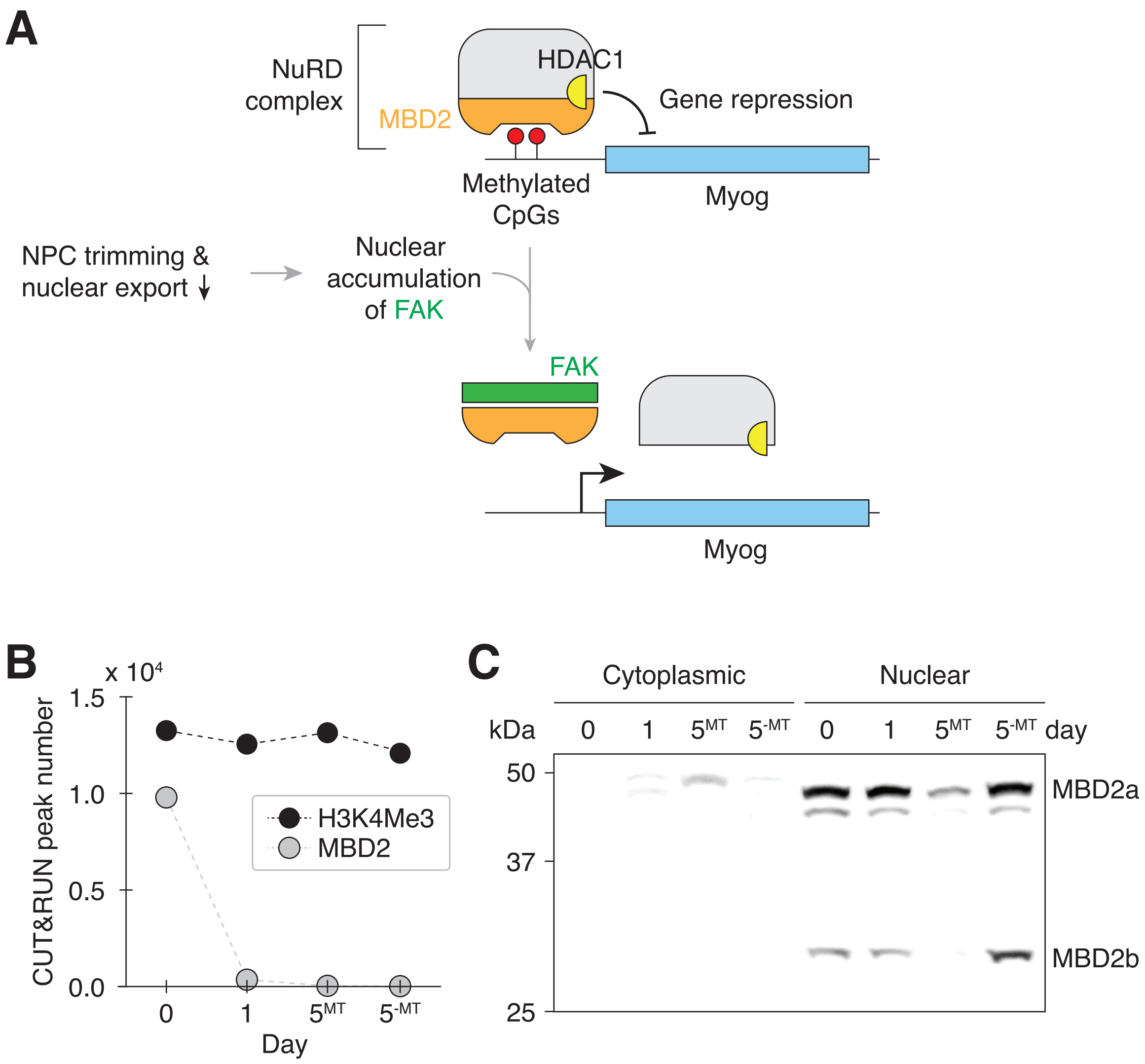


**Figure S9 I Transient nuclear accumulation of FAK resets MBD2-mediated genome regulation during myogenesis. (A)** NPC trimming is upstream of FAK-mediated MBD2 genome-binding reset event. **(B)** The number of MBD2 and H3K4Me3 CUT&RUN peaks in differentiating C2C12 cells. **(C)** Cytoplasmic and nuclear levels of MBD2 in differentiating C2C12 cells were determined by western blotting. 20 µg of protein was loaded per lane.

### Figure S10 (Ponceau S staining – Related to Figures 1, 2, 3, and 5)

Staining levels were determined by densitometric analysis. Normalized against lanes marked with arrowheads.

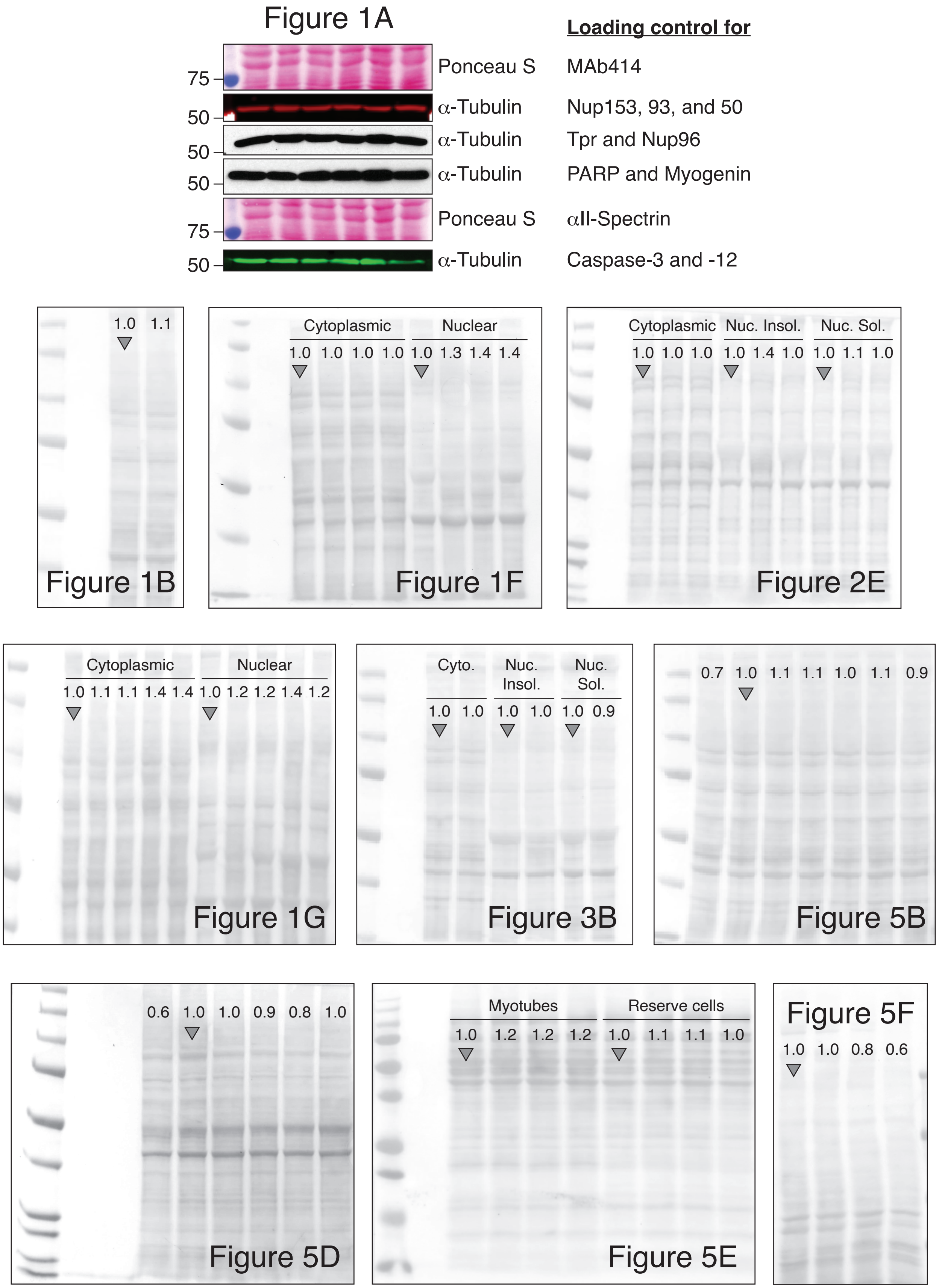

### **Figure S11** (Ponceau S staining – Related to Supplementary Figures)

Staining levels were determined by densitometric analysis. Normalized against lanes marked with arrowheads.

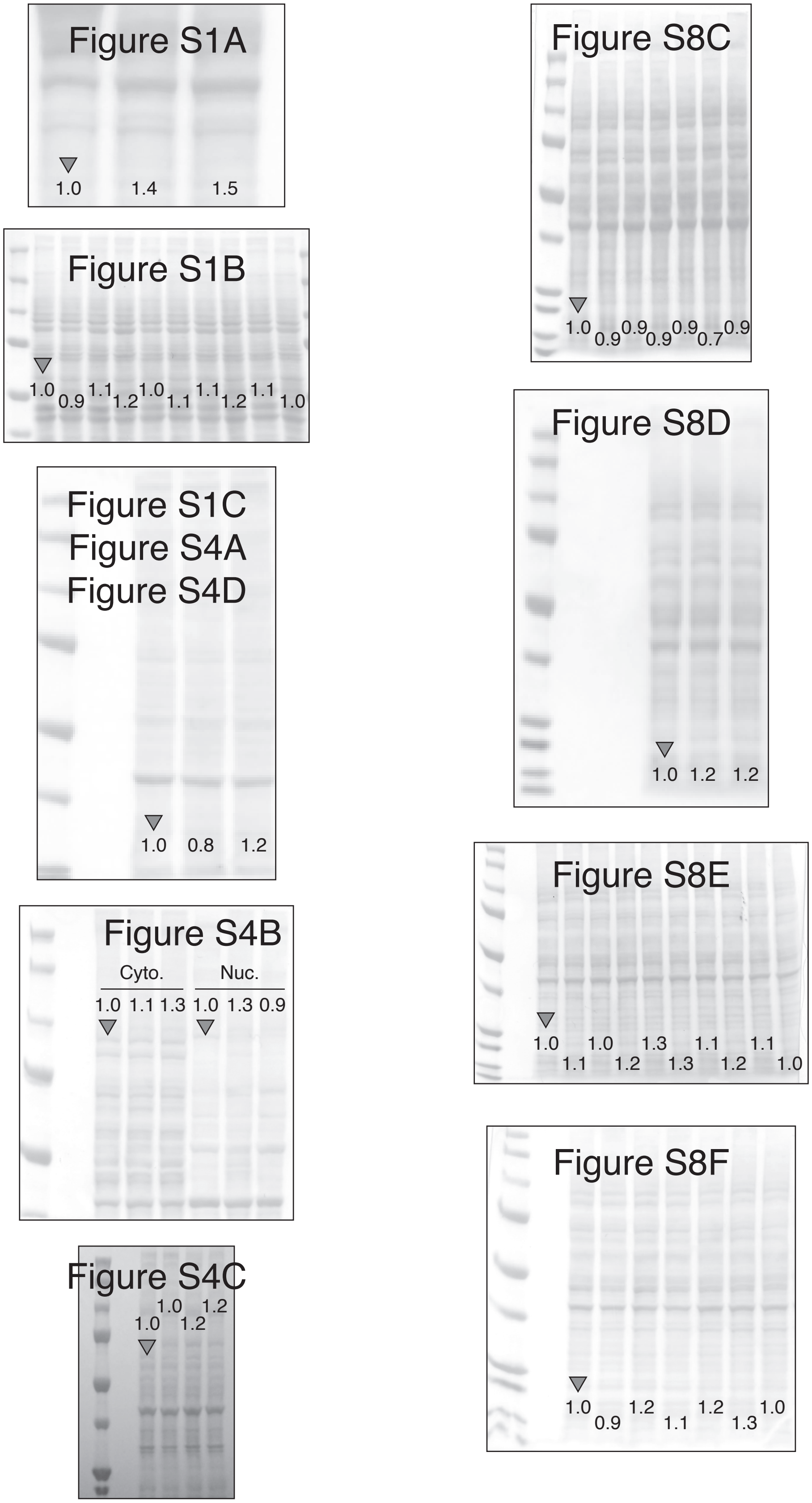
